# The Hox transcription factor Ubx ensures somatic myogenesis by suppressing the mesodermal master regulator Twist

**DOI:** 10.1101/2020.02.24.963231

**Authors:** Katrin Domsch, Julia Schröder, Matthias Janeschik, Christoph Schaub, Ingrid Lohmann

**Affiliations:** Heidelberg University; University Erlangen-Nurmberg

**Keywords:** Hox genes, Ultrabithorax (Ubx), mesoderm, twist, repression, Tinman, Tin, Mef2

## Abstract

Early determination factors and lineage-specific master regulators are essential for the specification of cell and tissue types. However, once a cell has committed to a specific fate, it is equally critical to restrict the activity of such factors to enable proper differentiation. In many studies the functional network for master regulators are under constant investigations. Yet, how these factors are silenced remains unclear. Using the *Drosophila* mesoderm as a model and a comparative genomic approach, we identified the Hox transcription factor (TF) Ultrabithorax (Ubx) to be critical for the repression of the mesodermal master regulator Twist (Twi). Mesoderm-specific Ubx loss-of-function experiments using CRISPR/Cas9 as well as overexpression experiments demonstrated that Ubx majorly impacts *twi* transcription. A detailed mechanistic analysis revealed that Ubx requires the function of the NK-homeodomain protein Tinman (Tin) but not the muscle differentiation factor Myocyte enhancer factor 2 (Mef2) to bind to the *twi* promoter. Furthermore, we found these TF interactions to be critical for silencing of the *twi* promoter region by recruiting the Polycomb DNA binding protein Pleiohomeotic (Pho). In sum, our study demonstrates that the Hox TF Ubx is a critical player in mediating the silencing of the mesodermal master regulator Twi, which is crucial for coordinated muscle differentiation.

## Introduction

The early development of an organism is tightly regulated by networks of transcription factors (TFs) that ensure axis orientation, germ layer determination and cell type specification as well as differentiation. Some of these TFs, so-called master regulators, play fundamental roles in cell and tissue specification, as they act at the top of hierarchical cascades driving cell fate decisions (Levine and Davidson, 2005). Thus, many studies focus on understanding the role of these master regulators during cell type specification (Jin et al., 2013; Liu et al., 2009; Sandmann et al., 2007). However, early master regulators lose their functional importance once cells have decided on their developmental path, and they need to be inactivated to enable proper cell and tissue differentiation (Davis and Rebay, 2017). Despite this knowledge, the mechanisms driving the inactivation of early master regulators, which is critical for tissue development, are not well understood.

The evolutionary conserved class of Hox TFs represent ideal candidates to coordinate late specification and differentiation events through the inactivation of master regulators for various reasons. First, they function after cell fates have been specified and they are active throughout the life time of an organism (Pearson et al., 2005). And second, they are essential for segment specification along the anterior-posterior axis and are expressed in all tissues within one segment. Thereby, they endow cells not only with a segment-specific code but also perform tissue- and cell-specific functions (Akam, 1987; Graham et al., 1989; Lewis, 1978; Scott and Carroll, 1987). In this study, we focussed on the Hox TF Ultrabithorax (Ubx), a Hox TF which is essential for the specification of the last thoracic and the abdominal segments (Lewis, 1978). Previous genome-wide investigation of tissue-specific Hox function identified Ubx as a repressor of alternative cell fates in the mesoderm, which required the cooperation with the Polycomb complex (Domsch et al., 2019). Thus, we assumed that Ubx could coordinate muscle differentiation by repressing not only alternative fate genes but also mesodermal master regulators.

Development of the *Drosophila* mesoderm depends on the nuclear translocation of dorsal (dl), a TF activating the mesodermal master regulator Twist (Twi) (Jiang et al., 1991; Thisse et al., 1991). Twi in turn induces the expression of additional cell-type specific factors, including the NK-homeodomain TF Tinman (Tin) and the MADS-Box TF Myocyte enhancer factor 2 (Mef2) (Cripps et al., 1998; Sandmann et al., 2007; Yin et al., 1997). Tin is the critical for visceral and dorsal somatic muscle specification as well as heart development, while Mef2 functions as a muscle differentiation factor (Azpiazu and Frasch, 1993; Bour et al., 1995). Due to its master-regulatory function, Twi has been extensively studied to elucidate its role in mesoderm determination and specification. Importantly, overexpression experiments revealed that Twi expression needs to be tightly regulated to prevent tissue malformation as well as loss and mis-location of cells (Baylies and Bate, 1996). In line, Twi expression is suppressed after mesoderm specification and is only maintained in adult muscle precursors (AMPs), a cell type that is silenced until remodelling of the muscle structure during metamorphosis (Bate et al., 1991). Thus, normal muscle differentiation requires the silencing of *twi* expression, however so far, the factors performing this critical function were unknown.

Here, we identified the Hox TF Ubx to inactivate the mesodermal master regulator Twi before muscle differentiation. Mesoderm-specific interference with Ubx maintained *twi* expression in many mesodermal cells and not just the AMPs, while premature activation of Ubx in the early mesoderm resulted in reduced *twi* expression. Importantly, we found that Ubx binds the *twi* promoter region together with the NK-homeodomain TF Tin and the MADS-Box TF Mef2, however required only Tin for this interaction. In addition, our results demonstrated that Tin and Mef2 are displaced from the promoter upon Ubx binding, which induced silencing of this region in cooperation with the Polycomb protein Pho. In sum, our study revealed that the Hox TF Ubx ensures normal muscle differentiation by repressing the mesodermal master regulator Twi.

## Results

### Ubx controls the expression of mesoderm specification and identity genes

We and others previously uncovered the Hox TF Ubx to activate mesodermal genes (Kremser et al., 1999; Manak et al., 1995) and to repress the expression of alternative fate genes, which we showed to be crucial for the stabilization of the mesodermal tissue lineage (Domsch et al., 2019). Re-analysing the transcriptome dataset uncovered that the Ubx-dependent control of tissue development is more complex than anticipated, as a substantial number of repressed Ubx targets were genes critical for mesoderm specification and differentiation. Based on our previous finding, we speculated that Ubx could control the repression of mesodermal identity genes by interacting with the Polycomb component Pleiohomeotic (Pho) to set repressive chromatin marks (Domsch et al., 2019). To test this assumption, we re-visited our established datasets (Domsch et al., 2019) and identified 504 chromatin regions bound by Ubx in the mesoderm that overlapped with active (H3K27ac) histone marks during mesoderm specification (stages 10-13, 4-9 hours after egg lay (AEL)), and switched to repressive marks (H3K27me3) during mesoderm differentiation (stages 14-17, 10-18 hours AEL) (Fig. 1B). As 88% of these genomic regions were also bound by Pho (Fig. 1C), these results indicated that these regions experience a Pho-dependent switch to inactivation. Gene Ontology (GO) analysis of the genes associated with these genomic regions identified a significant over-representation of terms linked to stem cell commitment, early mesoderm development and gastrulation (Fig.1B). Intriguingly, we found that histone marks and changes correlated with genomic location, as the majority of Ubx binding events that changed their epigenetic status were found at promoter regions (Fig. 1D,E, SupFig 1A). And again, genes associated with such promoter regions controlled early specification events of the mesoderm (mesoderm cell fate specification, stem cell commitment etc) and encoded preferentially TFs of different classes (Fig. 1D,E,F, SupFig 1A). Enhancer regions were associated primarily with late muscle functions and structure genes (Fig. 1D,E, SupFig 1A).

**Figure 1:**
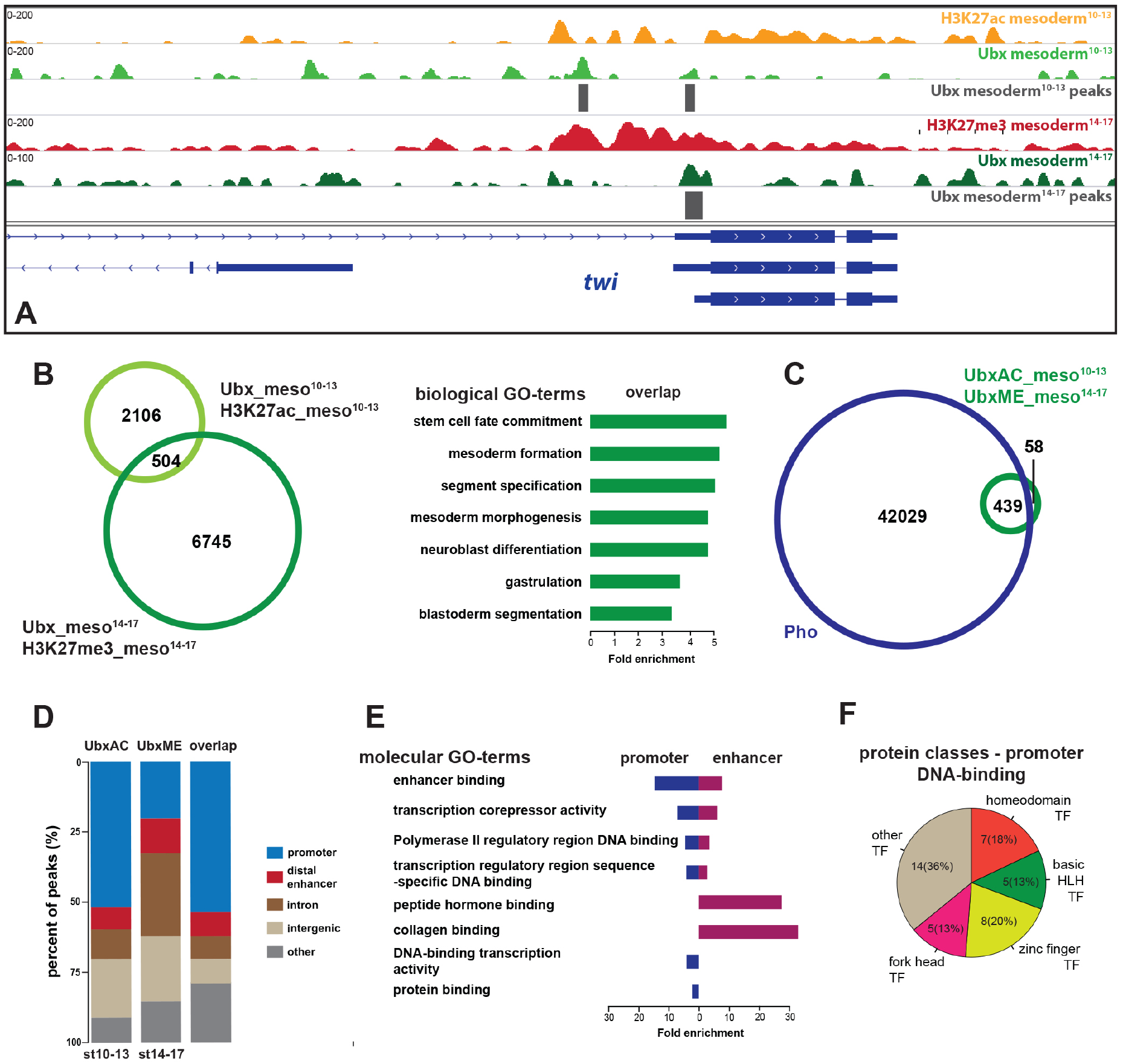
Ubx genomic interactions during mesoderm development. **A)** Illustration of the *twi* genomic region, showing Ubx binding and histone mark distribution at different developmental stages. Yellow: acetylation H3K27ac stages 10-13, red: methylation H3K27me3 stages 14-17, light green: Ubx stages 10-13, dark green: Ubx stages 14-17, grey boxes: accepted Ubx peaks, blue: coding region of the gene. **B)** Overlap of Ubx binding events associated with acetylated or methylated histone marks at different stages in the mesoderm, biological GO-term analysis (green) of the overlap. **C)** Overlap of Pho binding events and the genes co-bound by Ubx in different stages identified in B. **D)** Comparison of the localization of peaks called within unique genomic regions of Ubx. Localization are classified as promoters (−1000-+10 bp from TSS, 5’ UTR), distal enhancers (−2000 to −1000 from TSS, 3’ UTR, downstream), intron (intronic regions), intergenic (distal intergenic) and other regions (including exons). **E)** Molecular GO-term analysis of the overlap divided in enhancer (red) and promoter (blue) regions. **F)** Pie chart of the identified protein classes of genes that are bound by Ubx at the promoter region and have a function associated with DNA-binding.

The switch from active to repressive histone marks at promoter regions suggested that the associated genes get repressed in the course of mesoderm development. In order to provide direct evidence for this assumption, we tested whether the genes associated with these regions get de-repressed in the absence of Ubx. To this end, we overlapped the promoter-associated genes with genes previously identified as up-regulated when Ubx was mesoderm specifically degraded (Domsch et al., 2019). This analysis identified 32 genes, which fulfilled these requirements, including the early mesodermal master control gene *twist* (*twi*) (Fig. 1A). Importantly, *twi* expression, which persists only for a few hours throughout the mesoderm, gets induced very early during cellularisation to drive gastrulation and specification of mesodermal cell lineages (Leptin, 1991), while *twi* transcription is restricted to adult muscle precursors (AMPs) already in mid-embryogenesis (Bate et al., 1991).

In sum, these data showed that Ubx repressed the expression of mesoderm identity genes including *twi* and suggested that Ubx performed this function by controlling the establishment of repressive chromatin marks.

### Ubx is required for *twi* repression at the onset of mesoderm differentiation

In a next step, we investigated the requirements of Ubx in the mesodermal lineage in more detail. To efficiently eliminate Ubx activity in the mesoderm, we interfered with Ubx function using an alternative strategy to the protein degradation-based approach we had previously established, which strongly but not completely removed Ubx protein levels (Domsch et al., 2019). To this end, we targeted the *Ubx* gene using the inducible CRISPR/Cas9 mediated gene disruption, which relies on the tRNA-flanked multiplexed single guide RNA (gRNA) approach designed by Port and Bullock (Port et al., 2014; Port and Bullock, 2016) and the UAS/Gal4 system (Brand and Perrimon, 1993). Specifically, we generated a transgenic UAS line, which carries four Ubx specific gRNAs that bind within and close to the first Ubx exon, and combined it with the *UAS-Cas9.P2* transgene (SupFig. 2A). In addition, we also generated a CRISPR line for another *Hox* gene, *Antennapedia* (*Antp*), using the same strategy as for *Ubx* to validate the specificity of our gene targeting approach (SupFig. 2A).

First, we validated the efficiency of gene disruption by ubiquitously expressing *Ubx* or *Antp* gRNAs and Cas9 using the *armadillo* (*arm*)-GAL4 driver (Sanson et al., 1996). qPCR analysis of total RNA isolated from embryos of late developmental stages (stage 14-17) revealed that the gRNAs function gene specifically, as a significant reduction of *Antp* or *Ubx* transcript levels was only observed when the respective gRNAs were expressed (SupFig. 2B). Importantly, *Antp* transcript levels were increased when the *Ubx* gene was disrupted (SupFig. 2B), which is in accordance with the known cross-regulatory interaction of *Hox* genes (Morata and Kerridge, 1982; Struhl, 1982). Indeed, it has been shown before that *Antp* expression is expanded in *Ubx* mutants, leading to a change of the identity of abdominal segments A1 and A2 into the one of thoracic segments (Lewis, 1978).

After having confirmed the functionality and specificity of the *UbxCRISPR* system, we characterized the effects of interfering with *Ubx* specifically in the mesoderm in detail. Previously, we had used the *Mef2*-GAL4 driver, which starts to drive expression in embryonic stage 7 shortly before Ubx is active (Ranganayakulu et al., 1998). To interfere with the *Ubx* gene function already early in development, we combined the *twist-GAL4* (*2xPE twi-Gal4*) driver, which starts to be active at embryonic stage 5 (Baker and Schubiger, 1996), with the *Mef2-GAL4* driver. We investigated the effects of *UbxCRISPR* using this driver combination, which we refer to as *twi&Mef2-GAL4*, and performed immunostaining for Ubx and the muscle structure protein Tropomyosin 1 (Tm1). In comparison to control animals, Ubx expression in muscles was significantly reduced in *twi&Mef2>UbxCRISPR* embryos (Fig. 2A,A’,B,B’,E), while the neuronal expression was unaffected (Fig. 2C,D,E). In addition, the well-characterized Ubx expression in the visceral mesoderm, which marks the parasegment 7 (PS7), was absent in *twi&Mef2>UbxCRISPR* embryos (Fig. 2C,D, yellow arrow head). Loss of Ubx expression was accompanied with defects in morphology, as muscles were malformed, detached or missing in *UbxCRISPR* embryos, which was not the case in control embryos (Fig. 2A,A’’,B,B’’, highlighted with an yellow arrow head). This muscle pattern resembled in part the phenotypes obtained when maintaining *twi* expression using *Mef2-Gal4*, in particular longer muscles that are significantly thinner towards their ends when compared to wild-type embryos (Fig. 2A,A’’,B,B’’, SupFig. 2E-G). As *twi* was among the Ubx target genes that remained expressed in the absence of Ubx (Fig. 1, SupFig. 1), we hypothesized that one of the functions of Ubx during mesoderm development was the timed repression of master control genes like *twi* at the onset of mesoderm differentiation. To provide evidence for this, we performed *in situ* hybridisations to monitor *twi* expression in mesoderm-specific *Ubx* loss of function (LOF) embryos (mesodermal *UbxCRISPR*). Investigation of stage 14 embryos revealed that *twi* transcript levels were significantly increased in abdominal segments of Ubx LOF embryos when compared to control animals (Fig. 2F,G,H highlighted with the yellow arrows), while thoracic *twi* expression was unaffected (Fig. 2H). Conversely, when we induced Ubx expression already early during mesoderm specification using the *twi-GAL4* driver, *twi* transcript levels were reduced in particular in thoracic segments of stage 11 (but also of later stage) embryos (Fig. 2I,J,K, highlighted with the yellow arrows).

**Figure 2:**
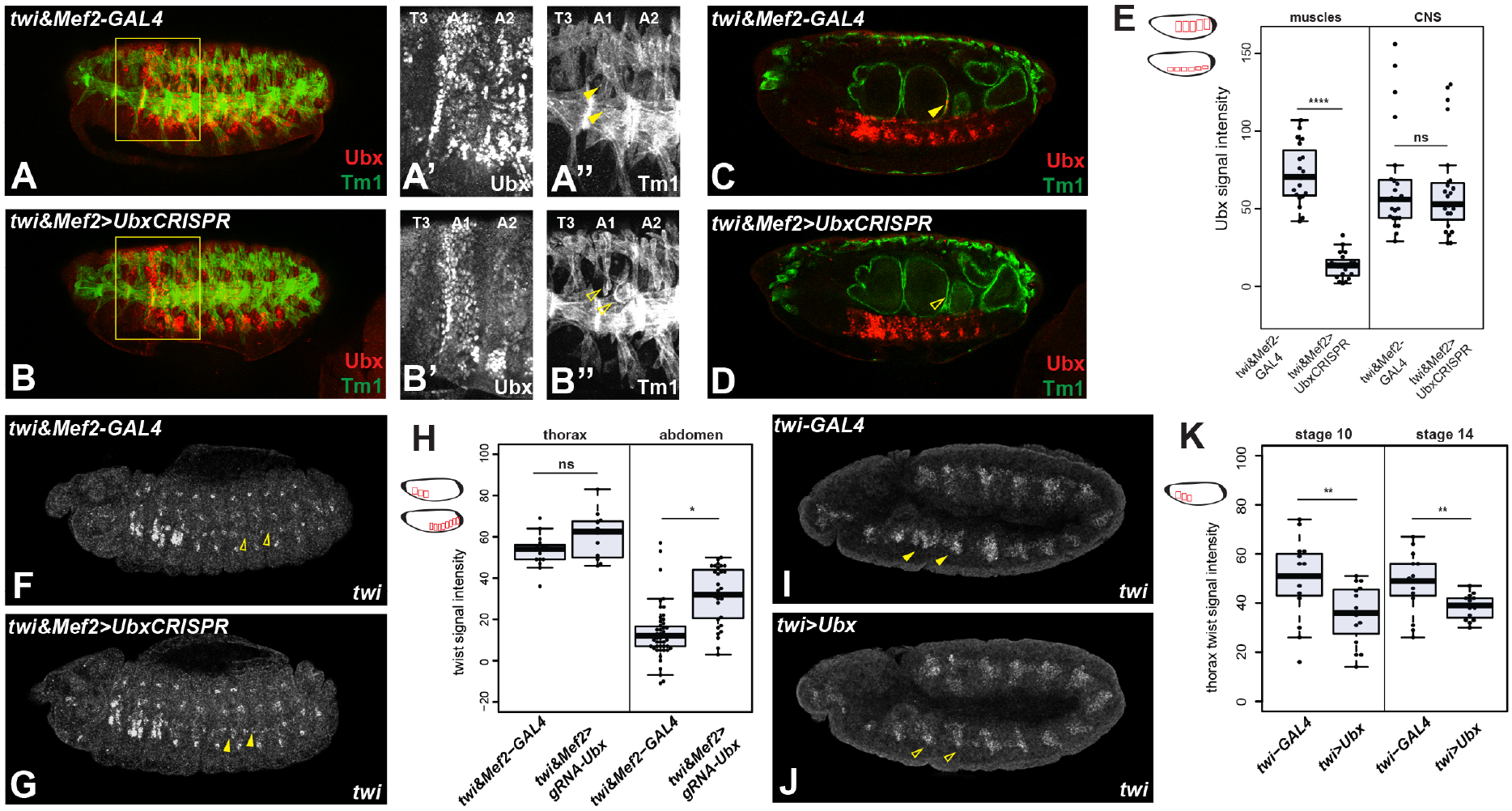
Ubx represses *twi* transcription. **A-D)** Control (*twi&Mef2-Gal4*) and *UbxCRISPR* (*twi&Mef2>UbxCRISPR*) stage 16 embryos stained with Tropomyosin (Tm1, green) and Ubx (red). Box highlights third thoracic and first, second abdominal segments. A’, B’) enlargement in grey scale, thoracic segment 3 (T3) and abdominal segments 1 & 2 (A1, A2), showing loss of Ubx expression in the mesoderm while expression in the ectoderm is unaffected. A’’,B’’) enlargement in grey scale of Tm1, open yellow arrowheads indicate the missing muscles in *UbxCRISPR* compared to the control (closed arrowheads). **C,D)** gut and CNS view of A and B, indicating the maintenance of Ubx expression in the CNS and missing visceral mesoderm expression in *UbxCRISPR* (empty arrow head) embryos. **E)** Validation of the Ubx expression in the muscle and CNS, illustration of the measured region in red, up: mesoderm, down: CNS, showing that the CNS expression is not affected by the *UbxCRISPR* but the mesodermal expression in decreased, **** p=3.02e-15. **F,G)** *in situ* hybridisation of control (*twi&Mef2-Gal4*) and *UbxCRISPR* (*twi&Mef2>UbxCRISPR*) embryos stage 14 using *twi* probe, closed arrowheads indicate additional *twi* expression in the *UbxCRISPR*. **H)** Validation of *twi* expression increase, illustration of the measured region in red, up: thorax, down: abdomen, showing that the thoracic expression is unchanged whereas the abdominal expression increased in *UbxCRISPR* experiments, * p=0.03, ns: non-significant. **I,J)** *in situ* hybridisation of control (*twi-Gal4*) and mesodermal Ubx overexpression (*twi>Ubx*) embryos stage 10 using *twi* probe, displaying a reduced *twi* expression in the overexpression (closed yellow arrow) **K)** Validation of *twi* expression decrease, illustration of the measured region in red (thorax), stage 10: ** p=0.01, stage 14: ** p=0.008.

In sum, these results highlighted that Ubx function was required and sufficient for *twi* transcriptional repression.

### Complex interplay of Ubx, Tin and Mef2 at the *twi* promoter

It has been shown previously that *twi* expression starts during cellularisation under the control of dl (Ray et al., 1991; Thisse et al., 1991) and maintains its own expression during mesoderm specification (Levine and Davidson, 2005; Sandmann et al., 2007). In addition, other TFs control *twi* expression (Sandmann et al., 2007), including Myocyte enhancer factor 2 (Mef2), which is generally required for muscle development and expressed in embryonic and larval muscles (Bour et al., 1995), and Tinman (Tin), which is restricted to the embryonic heart cell precursors by stage 12 (Azpiazu and Frasch, 1993) (Fig.3A). Importantly, Mef2 and Tin positively control *twi* transcription by interacting with the *twi* promoter (Sandmann et al., 2007), a region we found to be bound by Ubx (Fig 1A) (Domsch et al., 2019). This raised the intriguing possibility that Ubx replaces one or both of these TFs at the *twi* promoter thereby initiating *twi* repression at the onset of differentiation. To test this hypothesis, we studied the relationship of Ubx, Tin and Mef2 at the *twi* promoter. We performed ChIP-qPCR experiments for all three factors at three different time points, representing the determination and early specification (stages 7 to 9), the specification (stages 10 to 12) and the differentiation (stages 14 to 17) stages of the mesoderm (Fig. 3A). To precisely map TF interaction sites, we divided the *twi* promoter into four regions (Fig. 3A). We found Tin and Mef2 to strongly interact with all regions of the *twi* promoter at embryonic stages 7 to 9, while binding was significantly decreased during stages 10 to 12, which became even more pronounced during stages 14 to 17 (Fig.3B). The converse binding behaviour was observed for Ubx. Consistent with the late onset of Ubx expression, we did not detect Ubx interactions with the *twi* promoter at stages 7 to 9, while we found Ubx to specifically bind region 1 and 5 starting at stages 10 to 12, which was strongly increased at stages 14 to 17 (Fig. 3B). Intriguingly, region 5 experienced the most profound reduction in Tin and Mef2 binding as soon as Ubx started to interact with this region (Fig. 3B), suggesting that Ubx caused the clearance of Tin and Mef2 protein from the *twi* promoter. To test this possibility, we performed Tin and Mef2 ChIP-qPCR experiments in mesodermal cells devoid lacking *Ubx* gene expression (UbxCRISPR). We chose stage 10 to 12 embryos for this analysis, as this time frame represents the transition phase when *twi* expression starts to be repressed. Intriguingly, binding of both TFs was significantly increased at regions 4, 1 and 5 in the absence of Ubx (Fig.3C). As expression of Mef2 and Tin was unchanged in *Ubx* mutants (SupFig.3C,D,K), these results showed that Ubx plays an important role in preventing binding of these two TFs to the *twi* promoter.

**Figure 3:**
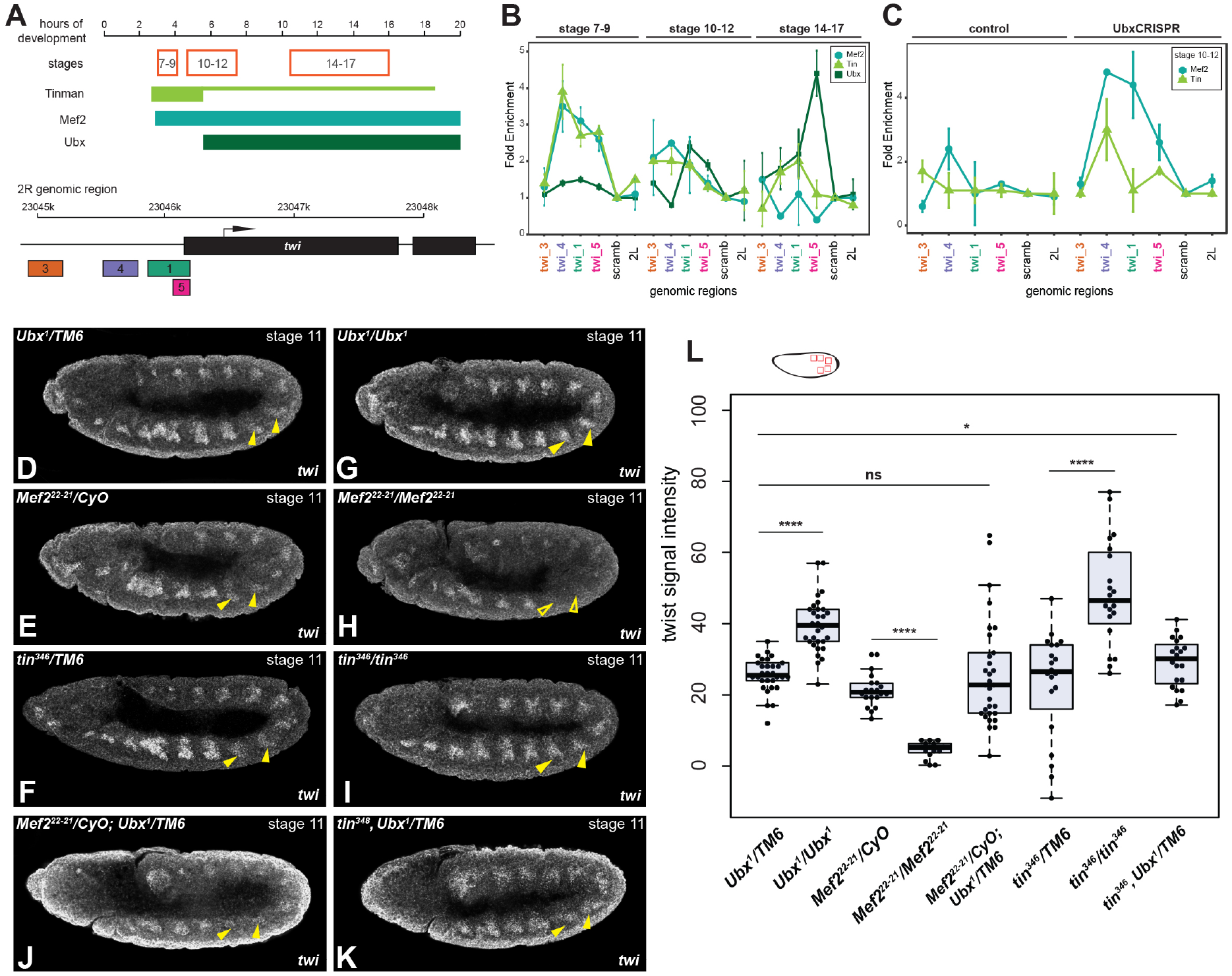
Ubx genetically interacts with Tin but not with Mef2. **A)** Up panel: time scale with analysed stages (red box), illustration of Tin, Mef2 and Ubx expression regarding the hours of the development. Lower panel: *twi* genomic region, illustration of the analysed promoter regions in a respected colour code. **B)** Mef2, Tin and Ubx ChIP-qPCR experiments on wildtype chromatin at different stages, showing that Tin and Mef2 bind to the *twi* promoter during stages 7-9 and then their binding is reduced in the following, whereas Ubx is expressed at stages 10-12 and starts to significantly accumulate at the promoter over time **C)** Mef2 and Tin ChIP-qPCR experiments using control (*twi&Mef2-Gal4*) and *UbxCRISPR* (*twi&Mef2>UbxCRISPR*) chromatin at stage 10-12, Mef2 and Tin binding is maintained in the Ubx know-down background. **D-K)** *in situ* hybridisation of *Ubx, tin* and *Mef2* heterozygous (D,E,F) and homozygous (G,H,I) mutants stage 11 using *twi* probe, yellow arrow heads indicate the abdominal segments 1 & 2, showing an increase of *twi* expression in *Ubx* and *tin* homozygous mutants (closed arrow) and decrease in *Mef2* mutants (open arrow), *in situ* hybridisation in double heterozygous background of *Mef2, Ubx* (J) and *tin, Ubx* (K) stage 11 show an increase of twi in *tin, Ubx* heterozygous conditions (closed arrow). **L)** Validation of *twi* expression, illustration of the measured region in red (abdomen), *Ubx*: **** p=1.03e-11, *Mef2*: **** p=3.68e-14, *tin*: **** p=3.56e-6, *tin, Ubx* heterozygous: * p=0.0467, ns: non-significant.

In a next step, we wanted to correlate TF interactions at the *twi* promoter with expression. Thus, we quantified *twi* transcript levels in abdominal segments of *Ubx*, *tin* as well as *Mef2* mutants. We found *twi* transcription to be unaffected in stage 11 *Ubx*, *Mef2* and *tin* heterozygous mutant embryos (Fig. 3D,E,F,L). Indeed, it resembled the characteristic *twi* expression in wild-type embryos with reduced *twi* transcript levels in abdominal compared to thoracic segments (SupFig. 2C,D). In contrast, *twi* transcription was increased in abdominal segments of *Ubx* and *tin* homozygous mutants (Fig.3G,I,L, highlighted by the yellow arrows), while expression was severely reduced in *Mef2* homozygous mutants (Fig.3H,L, highlighted by the yellow arrows). In addition, the expression of Ubx, Tin and Mef2 was unaffected in the respective mutant backgrounds (SupFig.3 A-D, I-K). This result indicated that Mef2 plays a critical role in maintaining the *twi* expression even at the onset of differentiation and not only during mesoderm specification. Furthermore, the correlation of increased Mef2 binding to the *twi* promoter (Fig. 3C) and increased *twi* transcription in the absence of Ubx (Fig. 3D,G,L) strengthened our hypothesis that Ubx replaces Mef2 at the *twi* promoter at the onset of mesoderm differentiation to inhibit *twi* activation. The interplay of Ubx and Tin is more complex, as binding of Tin to the *twi* promoter was also increased in the absence of Ubx (Fig. 3C). As for Mef2, this is indicative of a maintenance function of Tin, as *twi* transcription is increased in *Ubx* mutants (Fig. 2G,L). However, in contrast to Mef2 removal of Tin resulted in increased and not decreased *twi* expression levels (Fig. 3I,L), suggesting that Tin is directly or indirectly involved in the Ubx-mediated *twi* repression. As Ubx is expressed at similar levels in the mesoderm in *tin* mutants (SupFig.3 A,I), we assumed that Ubx requires Tin for the initial interaction with region 5 of the *twi* promoter, and not for later functions. In order to provide additional evidence, we investigated *twi* expression levels in heterozygous *Mef2, Ubx* as well as *tin, Ubx* double mutants. We found *twi* expression levels in heterozygous *Mef2, Ubx* double mutants to be comparable to single heterozygous conditions (Fig. 3J,L), whereas in *tin, Ubx* heterozygous conditions a significant increase of *twi* expression was noticeable.

These results indicated that Ubx interacts with Tin but not with Mef2 to initiate the repression of *twi*.

### Ubx requires Tin to interact with the *twi* promoter

To further investigate the requirements of Ubx in *twi* repression, we overexpressed *Ubx* in *Mef2* or *tin* homozygous mutants using the *twi-GAL4* driver and characterized *twi* transcript (and Ubx protein) expression in stage 10 embryos when Ubx is normally not active. Homozygous *tin* and *Mef2* mutant embryos did not display an obvious phenotype at this stage, as *twi* positive myoblasts were visible as a thin layer above the ectoderm (Fig.4A,C). In contrast, overexpression of Ubx in *Mef2* mutants resulted in a severe reduction of *twi* expression (Fig.4B), while Ubx overexpression in *tin* mutants did not affect *twi* expression (Fig.4D). These results demonstrated that Ubx required Tin function for *twi* repression.

**Figure 4:**
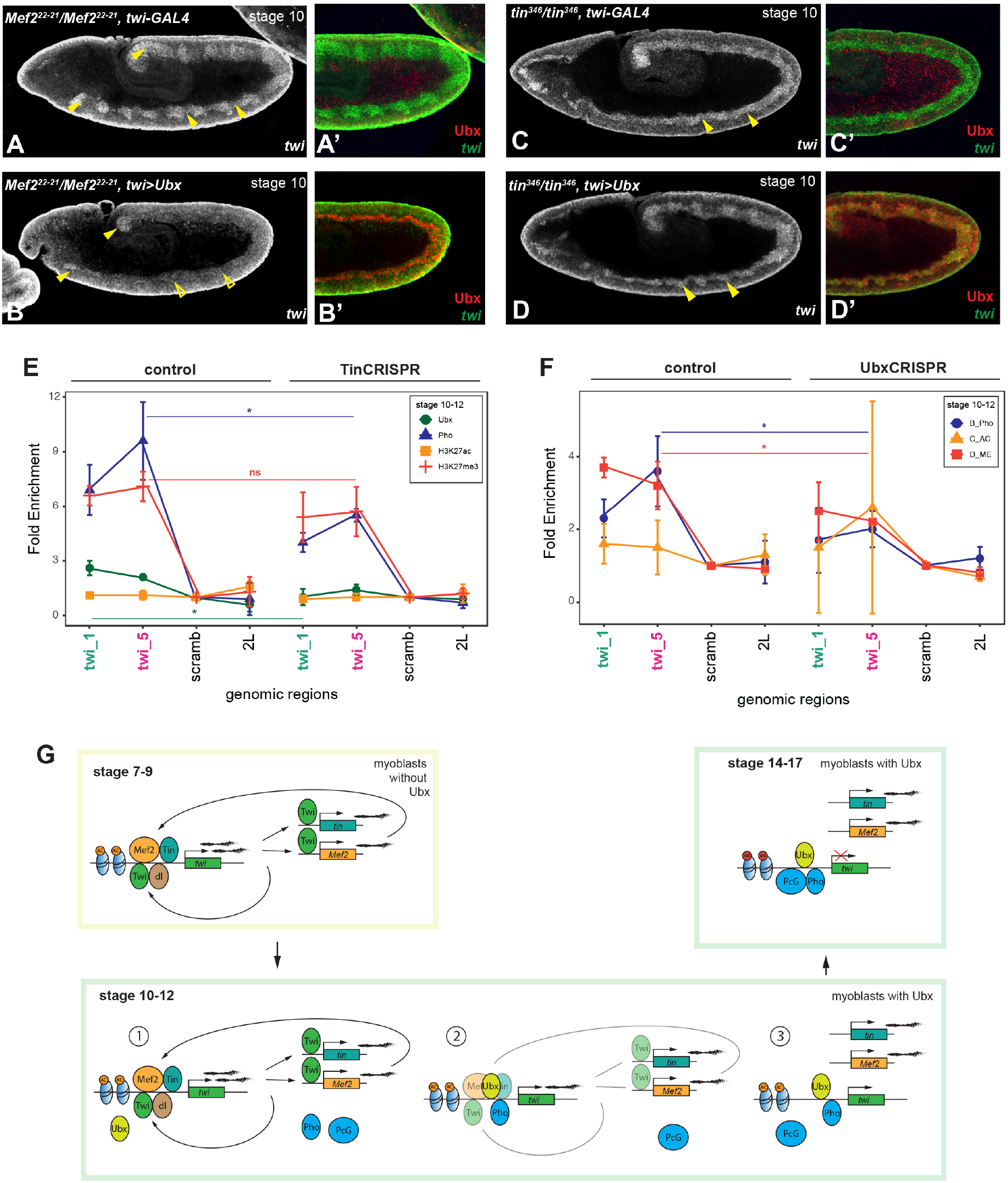
Ubx interaction with the *twi* promoter depends on Tin. **A-D)** *in situ* hybridisation of control homozygous *Mef2* and *tin* mutants containing the *twi-*GAL4 driver (A,C) and *Mef2 and tin* homozygous mutants (B,D) overexpressing Ubx stage 10 using the *twi* probe (white, green) and Ubx antibody (red) stainings, showing that the *twi* expression is reduced (open yellow arrows) in *Mef2* mutants overexpressing Ubx and is not affected in *tin* mutants overexpressing Ubx (closed yellow arrows). **E)** Ubx-ChIP (green), Pho-ChIP (blue), H3K27me3 (red) and H3K27ac (orange) experiments using control (*twi&Mef2-GAL4*) and *tinCRISPR* (*twi&Mef2>tinCRISPR*) chromatin at stage 10-12, indicating a decrease of Ubx (* p=0.011) and Pho (* p=0.029) binding to region 1 and 5 upon decreased Tin levels. H3K27me3 levels are decreased but not significant. H3K27ac levels are unaffected. **F)** Pho-ChIP (blue), H3K27me3 (red) and H3K27ac (orange) experiments using control (*twi&Mef2-GAL4*) and *UbxCRISPR* (*twi&Mef2>UbxCRISPR*) chromatin at stage 10-12, indicating a decrease of Pho (* p=0.0230) binding and H3K27me3 levels (* p=0.050) to region 1 and 5 upon decreased Ubx levels. H3K27ac levels are slightly but not significantly increase. **G)** Predicted model, stage 7- 9: Mef2 and Tin bind to the *twi* promoter and maintain its activity, stage 10-12: Ubx starts to be expressed in the mesoderm and replace Mef2, interacts with Tin and attracts Pho, the expression of *twi* gets reduced, stage 14-17: Ubx interact with Pho and attract PcG to maintain the inactivation.

In a next step, we analysed the interdependency of Ubx and Tin in more detail by testing the binding of Ubx to the *twi* promoter in the absence of Tin. To this end, we abolished Tin function in them mesoderm by using the inducible CRISPR/Cas9 mediated gene disruption to target the *tin* gene (SupFig. 3E-H). Mesoderm-specific expression of a *tin* specific gRNA (*tinCRISPR*) by means of the *twi&Mef2-GAL4* driver resulted in a significant reduction of Tin protein levels in *twi&Mef2>tinCRISPR* embryos in comparison to control (*twi&Mef2-Gal4*) embryos at stage 11 (SupFig. 3 F-H), validating efficient *tin* gene disruption. Subsequently, we performed Ubx-ChIP analysis in *tinCRISPR* embryos focusing on the *twi* promoter regions 1 and 5, which revealed a significant reduction of Ubx binding to these sites in the absence of Tin (Fig.4E). We had shown previously that Ubx repressed the expression of alternative fate genes by stabilising binding of the Polycomb protein Pho and the setting of repressive chromatin marks at control regions close to such genes (Domsch et al., 2019). Thus, we tested whether a similar mechanism applied to the repression of the master control gene *twi*, and investigated Pho binding, H3K27me3 and H3K27ac levels at the *twi* promoter in the absence of Tin and Ubx. Using ChIP-qPCR, we found Pho binding and H3K27me3 levels to be significantly reduced when the *tin* or the *Ubx* gene were disrupted, while H3K27ac levels were not changed in comparison to the control (Fig.4E, F). The results indicated that Ubx employs a similar mechanism as for the alternative fate gene repression and interacts with Pho at the *twi* promoter region. Further on, Ubx requires Tin also at the molecular level to interact with the *twi* regulatory element.

## Discussion

Master regulators instruct cells to adopt specific fates by activating complex gene regulatory networks. One such master regulator is Twi, which is critically required for mesoderm determination and specification in *Drosophila*. In order to progress in development and to enable differentiation, early specification factors need to be silenced, which is also the case for Twi during mesoderm development. However so far, the factors performing this critical function are largely unknown. Our study now identified the Hox TF Ubx to be critical for the inactivation of the mesodermal master regulator Twi, thereby ensuring the transition from mesoderm specification to differentiation. Based on our data, we propose the following model (Fig.4G). During late stages in mesoderm determination (stages 7 to 9), the *twi* promoter is bound by Tin and Mef2 to maintain *twi* expression. At the beginning of specification (stages 10 to 12), both factors are replaced by Ubx, which recruits the Polycomb protein Pho and initiates repression of *twi* transcription by establishing repressive chromatin marks. Interaction of Ubx with the *twi* promoter for repression requires Tin, while Mef2 seems to ensure the proper timing of *twi* activation versus repression. Once *twi* repression is mediated under the control of Ubx and Pho, coordinated differentiation of specified myoblasts into functional muscle fibers can be initiated (stages 14-17).

How master-regulatory networks are changed for the transition into the next developmental stage has been studied in a few cases, which led to the formulation of two mechanisms: 1) a context-specific network rewiring, and/or 2) a switch of regulatory relationships to assemble new control hierarchies (Atkins et al., 2013; Davis and Rebay, 2017).

During *Drosophila* eye development, a positive feedback loop essential to maintain Eyeless (Ey) expression is disrupted to proceed with photoreceptor fate specification and neuronal differentiation (Atkins et al., 2013). The inactivation of *ey* is achieved by unwiring the network and promoting the interaction of Eyes absent (Eya) and Sine oculis (So), which leads to direct repression of *ey* in differentiating cells and breaks the Ey regulatory network (Atkins et al., 2013). In our case, the Twi network is essential for mesoderm determination and specification, since Twi activates essential lineage identity genes that help to maintain its activation (Sandmann et al., 2007). The need for a tight regulation of *twi* expression is indicated by overexpression experiments showing severe tissue malformations (Baylies and Bate, 1996) and muscle abnormalities by maintaining Twi activity (SupFig.2). We assume the binding of Ubx to the *twi* promoter to mark the transition from specification to differentiation by downregulating *twi* and inducing a break/unwiring of the Twi network (Fig4). Repression of *twi* uncouples the Tin and Mef2 networks, which maintain an independent activity within the developing heart precursors as well as in myoblasts that will differentiate into functional muscle fibers, respectively (Azpiazu and Frasch, 1993; Bour et al., 1995; Sandmann et al., 2007).

Hox TFs are themselves master regulators of segment identity (Pearson et al., 2005). For example, loss of Ubx leads to a well-known homeotic transformation of the abdominal into thoracic segments promoted by Antennapedia (Antp), which shows that these TFs initiate their own regulatory network (Lewis, 1978). In the mesoderm, the inactivation of *twi* may lead to the activation of the Ubx control network. This network will in turn promote muscle differentiation and stable lineage commitment through the activation of genes for the progression of tissue development, like *decapentaplegic (dpp)* and *betaTubulin 60D (betaTub60D)* in the visceral mesoderm (Kremser et al., 1999; Manak et al., 1995). Importantly, we assume differentiation process to not inhibited but negatively influenced, since maintaining the *twi* expression during muscle differentiation leads to malformations but not severe muscle pattern or structure defects. Twi itself activates the muscle differentiation factor Mef2. Intriguingly, it is known that increased amounts of Mef2 proteins have a drastic impact on normal muscle formations and structures (Huang, 2000; Molkentin and Olson, 1996). This indicates that the phenotype observed in mild *twi* overexpression or loss of Ubx function experiments is caused by increased expression of Mef2, resulting in an imbalance in protein amounts (Huang, 2000; Molkentin and Olson, 1996).

To switch or unwire the Twi network, Ubx has to interact with the regulatory regions of *twi*. Our data, which show that Pho is unable to bind to the *twi* promoter in the absence of Ubx, suggest that Ubx binds to the *twi* promoter and recruits Pho to initiate *twi* inactivation via the assembly of the Polycomb complex. In addition, we find that Ubx is unable to interact with this region without Tin, suggesting that this TF “prepares” the chromatin for Ubx binding. This is in line with previous studies showing Tin serves as a chromatin anchoring protein in ventral myoblasts (Busser et al., 2013; Liu et al., 2009).

The function of Mef2 in *twi* repression is less clear, since the expression of *twi* is unaffected in *Mef2* mutants. However, our data also suggest that Mef2 plays a role, as the Ubx mediated repression of *twi* in the abdominal segments occurs earlier in the absence of Mef2. Thus, we assume Mef2 to control the timing of specification and differentiation. It has been shown before that Mef2 protein binds to the minor groove and bends the DNA in cooperation with co-factors (Huang, 2000). Taken together, we assume the Twi regulatory feedback loop is unwired through the binding of Ubx to the *twi* promoter region (Fig.4J).

In future, it will be important to also focus on the inactivation of essential master regulators and how the repression complex is established and formed, to understand in more detail how cells, which are devoted for differentiation, enter their developmental path.

## Materials and Methods

### Fly stocks and husbandry

For CRISPR experiments, the following lines were used: *UAS-Cas9.P2* obtained from Filip Port, *Mef2-GAL4* (BL50742), *twi-(2xPE)*-*GAL4* (BL2517), *arm-GAL4* (BL1561), *w*^1118^ (BL3605), *Mef2^22-21^/CyO wg-lacZ* given by Hanh Nguyen, *tin^346^/TM3 ftz-lacZ* gift from Manfred Frasch and *Ubx^1^/TM6 Dfd-LacZ* from Ana Rogulia-Ortmann, *UAS-gRNA-Ubx, UAS-gRNA- Antp*, *UAS-gRNA-Tin* and *UAS-HA-Ubx* overexpression constructs were made during the study. The mutants and overexpression constructs were crossed together accordingly for genetic interaction experiments.

### Generation of Ubx, Antp & tin UAS-gRNA and UAS-HA-Ubx constructs

For the UAS-gRNAs: The gRNAs were design using the technique and tool published by Filip Port implying tRNAs separated target-specific gRNAs (F. Port et al., 2014; Port and Bullock, 2016)in combination with the UAS/GAL4 system (Brand and Perrimon, 1993). Suitable gRNAs were identified using the webtool CRISPR Optimal Traget Finder (http://targetfinder.flycrispr.neuro.brown.edu) and the first exon as well as including flanking regions intronic regions for *Ubx* and *Antp*. The 20 nt were amplified and cloned in the pCFD6 plasmid according to the protocol published on https://www.crisprflydesign.org (Ubx-gRNA1: GGAGGCGTACAGAGCGGCGT, Ubx-gRNA2: GGCGAGCGGTAAAGCGCTGA, Ubx-gRNA3: GGGCGCCGCTGCCCAAACGG, Ubx-gRNA4: AAGAGTAGGTCAGCCGAAGG, Antp-gRNA1: GATGACGCTGCCCCATCACA, Antp-gRNA2: GGCCGTTGTAGTAGGGCATG, Antp-grNA3: GGCGGGATCAGCAGACGCTG, Antp-gRNA4: GGTTCTGATGGACCTGTGAT, Tin-gRNA1: GAGCATCTACGGTTCAGATG, Tin-gRNA2: GGACCTCAACAGCTCCGCTG, Tin-gRNA3: GCTTCTGCGTCGGAGCTTGC, Tin-gRNA4: GAGGGCGTGACGTATGGCGT).

For the overexpression construct: The full Ubx coding region (Ubx isoform I) was cloned into the pUAS-attB vector (Bischof et al., 2007) using a forward primer containing a EcoRI restriction site and the HA tag sequence as well as a revers primer with a XbaI restriction site. The UAS-constructs were injected by BestGene on attP5 (second chromosome, gRNA, UAS-Ubx) and attP2 (third chromosome, gRNA) carrying fly lines. Primers and sequence maps are available upon request.

### ChIP experiments and isolation of total RNA

ChIP experiments were performed as described in Sandmann et al., 2007 and analyzed by using qPCR experiments (Invitrogen Syber-Green-Mix, Primers are available upon request). The following antibodies were used: H3K27ac (ab4729, Abcam), H3K27me3 (ab6002, Abcam), gp-Ubx (Domsch et al., 2019), Rb-Mef2 (gift from Hanh Nguyen) and Rb-Tin (gift from Manfred Frasch (Jin et al., 2013). ChIP-qPCR results were calculated as described in Schaub et al., 2012. Total RNA was isolated with TRIZol (Invitrogen), used to generate cDNA (TermoFisher Scientific, RevertAid First Strand cDNA Synthesis Kit) and the material was analysed or validated by qPCR (Invitrogen Syber-Green-Mix, ChIP-Primers: *twi* promoter region 1 (twi_1): for GCCGACAATTCCCCTCGTAT, rev GGCGGGGGATTTCGATTTTC; *twi* promoter region 3 (twi_3): TTTTCGTGGAGTTCCCCTCG, rev TGGGATCGCAGGAATTTCCC; *twi* promoter region 4 (twi_4): for CGCTGCAAACAACAACATTCA, rev TCTTTGGATCGACGGGAATTCA; *twi* promoter region 5 (twi_5): for CGTCGAGTCAAGGCTCTCTT, rev CATCCCGCTCCCACTCAATG; *scramb1* (negative control): for GAGCGAAAAGGTGCAGAAAG, rev CCTGTCTGTTTCCCTGCTGT; region on the 2^nd^ chromosome left arm *2L* (negative control): for AGGTGTTGTTGTGGGTCCTT, rev TCCCAGAGTTCCCTTTAGCA; RNA-Primers: Ubx-Primer: for AGTTTACACCAGGCCAG CAA, rev GCCCGCAAGAGATTCTGAGT; Antp-Primer: for GGGCTATACGGACGTTGG AG, rev ATGTCCCTGGTGCATCTGTG; RpL32-Primer (Steiner et al., 2012): for TACAGGCCCAAGATCGTG AA, rev TCTCCTTGCGCTTCTTGGA).

### Bioinformatics analysis

Bioinformatics analysis was performed as described in (Domsch et al., 2019). Material and files used for the analysis were generated and published in (Domsch et al., 2019) (storage: NCBI Gene Expression Omnibus GSE121754, GSE121670, GSE12175). Visualization of the data was performed as described in (Domsch et al., 2019).

### Immunofluorescence staining and microscopy

Embryonic stainings were performed as described in (Domsch et al., 2019) and the following antibodies were used: Rat-Tm1 (1:200, MAC-141, ab50567, Abcam), gp-Ubx (Domsch et al., 2019), Rb-Tin (1:1000, gift from Manfred Frasch, (Jin et al., 2013)) and gp-Twi (1:500, gift from Eileen Furlong). For in-situ hybridizations a 1000 bp fragment of the *twi* was amplified and served as a probe template. The generation of the probe and the in-situ hybridization was performed as described in (Schindelin et al., 2012). Amplification was obtained with the TSA system (Perkin-Elmer). Biotinylated (1:200, Vector Labs) and fluorescent (1:200, Jackson ImmunoResearch or Invitrogen) secondary antibodies were used. Images were acquired on the Leica SP8 Microscope (20x/0.7 HC PL APO Glycerol, 63x/1.3 HC PL APO Glycerol). The collected images were analyzed and processed with Leica Application Suite X (LAS X, Leica Microsystems) and Adobe Photoshop.

### Quantifications of data

Quantification of the GFP signal intensity: GFP signal was measured with the FIJI/ImageJ (v1.52p) (Schindelin et al., 2012) from at least 5 segments per embryo in 5 embryos. The mean and the t-test for significance were calculated in Microsoft Excel and graphic illustrations were performed in R (R Core Team, 2018).

## Acknowledgments

We are grateful and like to thank the fly community and especially Filip Port for CRISPR fly stocks, Hanh Nguyen, Manfred Frasch, Ana Rogulia-Ortmann and Eileen Furlong for antibodies as well as BestGene and the Lohmann lab for critical comments on the manuscript. The project was founded by the DFG grand LO 844 8/1 to Ingrid Lohmann.

## Authors contribution

Conceptualization: K.D.

Methodology: K.D., J.S., M.J., C.S.

Validation: K.D., J.S.

Investigation: K.D., J.S., M.J.

Data curation: K.D.

Visualisation: K.D., J.S.

Supervision: K.D.

Project administration: K.D., I.L.

Funding acquisition: I.L.

Writing – original draft: K.D., I.L.

Writing – review & editing: K.D., I.L.

---

**SupFigure 1:**
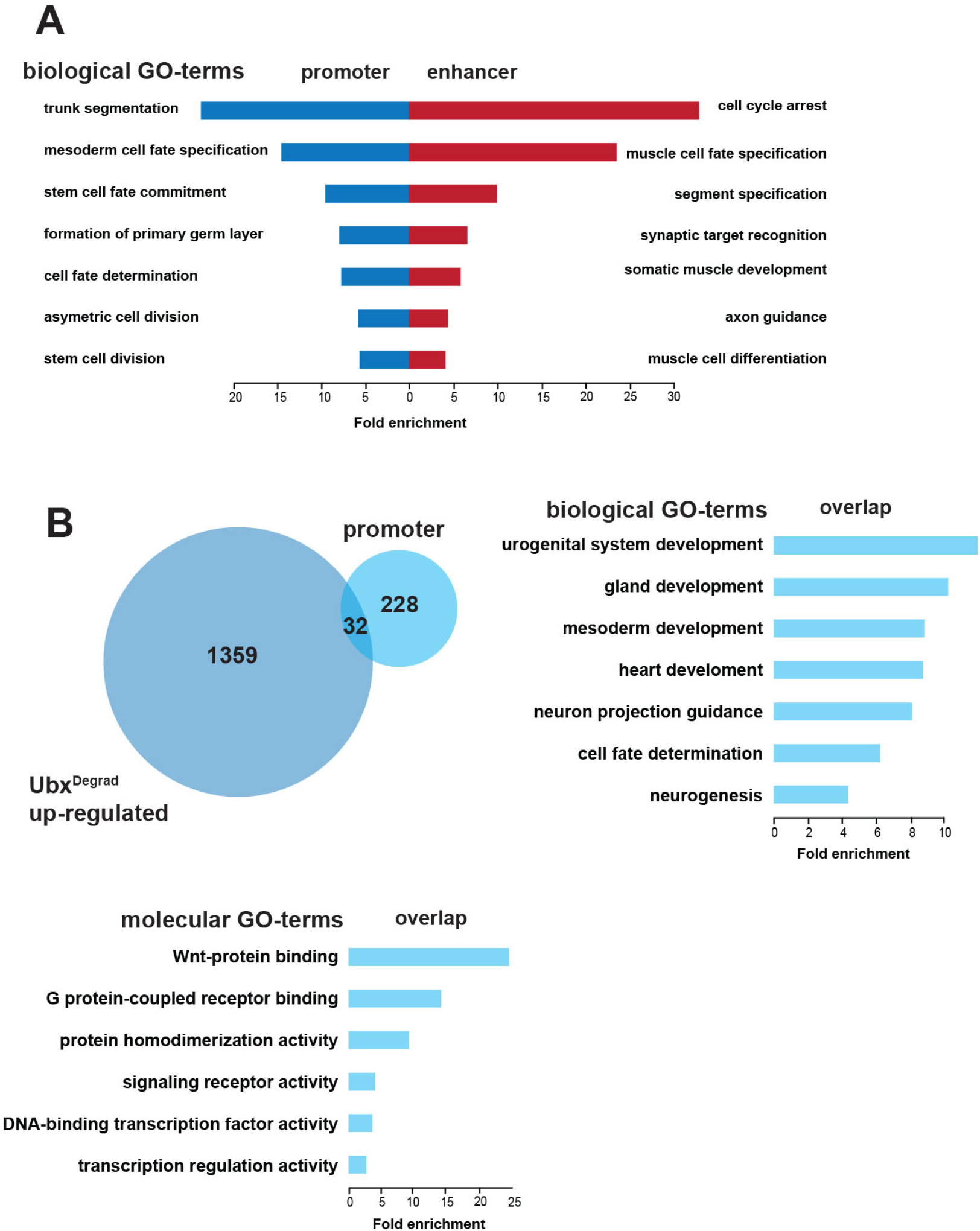
Ubx genomic interactions during mesoderm development. **A)** Biological GO-term analysis of the overlap from Fig.1B divided in enhancer (red) and promoter (blue) regions. **B)** Overlap of genes that are up-regulated in the UbxDegrad (dark blue) background and promoter regions from the Ubx overlap shown in Fig.1B (light blue), biological and molecular GO-term analysis of the 32 genes from the overlap in B.

**SupFigure 2:**
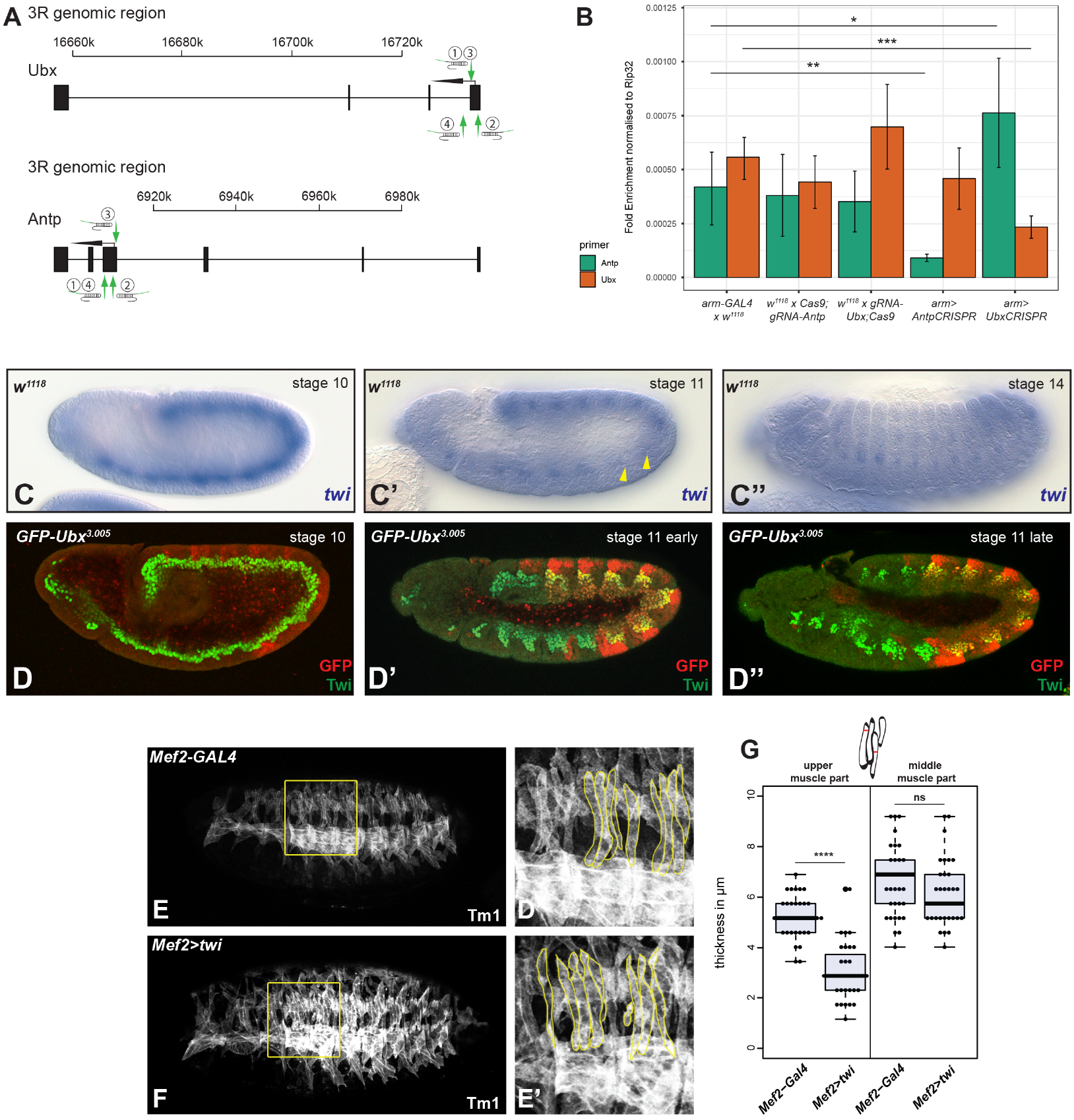
Ubx represses the *twi* transcription. **A)** Localisation of the *Ubx* and *Antp* gRNA. **B)** Validation of the *Ubx*- (orange) and *AntpCRISPR* (green) using *arm-Gal4*. qPCR results show a downregulation of *Antp* transcripts in the *AntpCRISPR* and *Ubx* transcripts in the *UbxCRISPR* when compared to the controls. In addition, *Antp* transcripts are upregulated in the *UbxCRISPR. Ubx in UbxCRISPR*: *** p=0.00074, *Antp in AntpCRISPR* ** p=0.0137, *Antp in UbxCRISPR*: * p=0.0089. **C)** *insitu* hybridisation of *w^1118^* embryos (wildtype) of different stages using *twi* probe, closed yellow arrow indicates the abdominal segments A1 und A2. **D)** Endogenous GFP-Ubx embryos stained with GFP (red, labelling GFP-Ubx) and Twi (green) at different stages. **E**,**F)** Control (*Mef2-Gal4*) and *twi* overexpression experiments (*Mef2>twi*) stage 16 embryos stained with Tropomyosin (Tm1), illustrating the lateral muscle pattern in. Box highlights third thoracic and first, second abdominal segments. **E’,F’)** Enlargement and highlighted (yellow) lateral transversal and segment boundary muscles. **G)** Validation of the muscle thickness lateral transversal muscles, illustration of the measured region above and measured region indicated in a red line, showing that the middle thickness is not affected but the end of the muscles show significant defects in thickness, **** p=1.10e-10, ns: non-significant.

**SupFigure 3:**
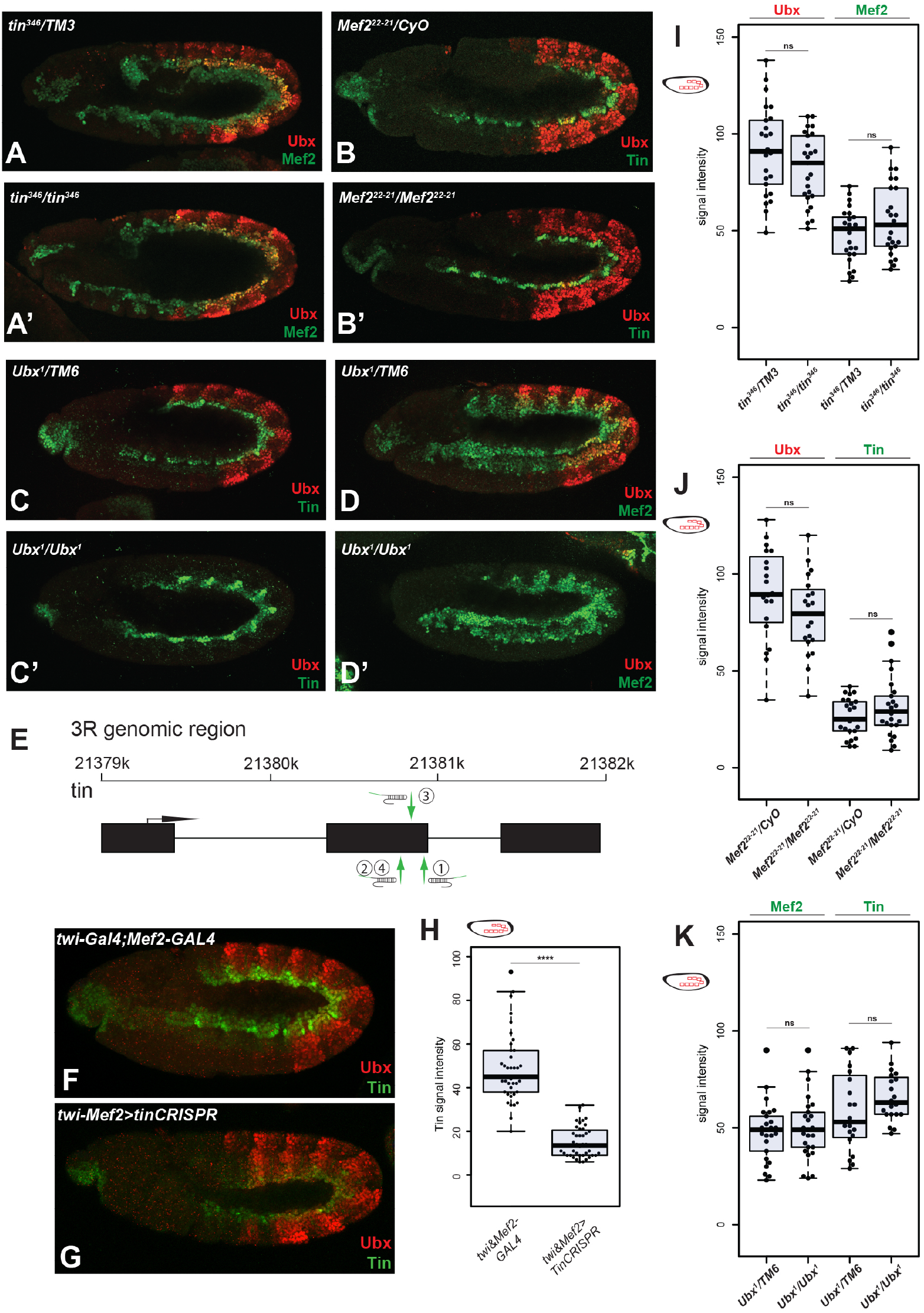
Ubx genetically interacts with Tin but not with Mef2. **A-D)** Ubx, Tin and Mef2 mutants stage 11, stained with Ubx (red), Mef2 (green) and Tin (green). **E)** Localisation of the *tin* gRNA. **F,G)** Control (*twi&Mef2-Gal4*) and *tinCRISPR* (*twi&Mef2>tinCRISPR*) embryos stage 12 stained with Ubx (red) and Tin (green) antibodies, showing an decrease in Tin expression whereas Ubx expression remains. **H)** Validation of decreased Tin expression, illustration of the measured region in red (abdomen), **** p=2.50e-18. **I,J,K)** Validation of the protein expression from A-D indicated at the top of each panel, illustration of the measured region in red (abdomen) focusing on the somatic myoblasts, ns: non-significant.

